# *Porphyromonas gingivalis* promotes lipid droplet-mediated microglial dysfunction

**DOI:** 10.64898/2026.05.03.722306

**Authors:** Muhammad Shahid Riaz Rajoka, Kristina Nicole Valladares, Chloe La Prairie, Wei Li, Peter King, Jannet Katz, Suzanne M. Michalek, Ping Zhang

## Abstract

Growing evidence supports a strong association between periodontitis and Alzheimer’s disease (AD), yet the mechanisms linking these conditions remain poorly defined. In neurodegenerative disorders, including AD, microglia are often characterized by increased lipid droplet (LD) accumulation, heightened activation, and impaired function. In this study, we examined whether *Porphyromonas gingivalis* (*Pg*), a keystone periodontal pathogen, promotes LD accumulation in microglia and disrupts their function. We found that *Pg* infection induces robust LD accumulation in BV2 microglial cells and in microglia from *Pg*-infected *App* KI mice. This *Pg*-driven LD buildup was closely associated with elevated reactive oxygen species (ROS) production, impaired phagocytic ability, and altered activation. Notably, pharmacological inhibition of LD with a triglyceride synthesis inhibitor effectively reversed *Pg*-induced LD accumulation, mitigated ROS production, and restored phagocytic function, thus underscoring the critical role of lipid metabolism in regulating microglial function. These findings support a model in which, in the context of periodontitis, systemic dissemination of periodontal pathogens promotes LD accumulation in microglia, and this metabolic alteration exacerbates microglia dysfunction via a self-reinforcing cycle of excessive oxidative stress and impaired phagocytosis, potentially accelerating AD progression.

## Introduction

Alzheimer’s disease (AD) is the most common cause of dementia, characterized by an age-related, progressive, and irreversible decline in cognitive and behavioral functions within the central nervous system (CNS) (Klyucherev et al. 2022). AD and its related dementias not only severely reduce patients quality of life but also bring a heavy economic burden to their families and society (Cavaco et al. 2025). The pathological hallmarks of AD include extracellular amyloid-β (Aβ) plaques and intracellular neurofibrillary tangles composed of hyperphosphorylated tau (Knopman et al. 2021). Aβ accumulation can be detected long before clinical symptoms appear, with its aggregation triggering a cascade of pathogenic events contributing to disease progression (Uhlmann et al. 2020). Despite extensive research, the mechanisms underlying AD remain incompletely understood, and no curative therapy is available (Passeri et al. 2022). Therefore, identifying modifiable risk factors is crucial to developing strategies to delay, mitigate, or prevent disease onset and progression.

Epidemiological studies have indicated multiple AD risk factors, including aging, systemic inflammation, traumatic CNS injury, and infection (Heneka et al. 2015). Periodontitis is a chronic inflammatory disease of the periodontium driven by dysbiosis of subgingival biofilm and is the leading cause of tooth loss in adults (Lamont and Hajishengallis 2015). Notably, the keystone periodontal pathogen *Porphyromonas gingivalis* (*Pg*) and its virulence factors have been detected in AD brains and linked to elevated Aβ and tau pathology (Dominy et al. 2019). Furthermore, patients with elevated serum antibody titers against *Pg* are associated with elevated cerebrospinal fluid levels of both Aβ and tau (Laugisch et al. 2018). Animal studies further demonstrate that *Pg* and its LPS promote Aβ deposition, tau hyperphosphorylation, neuroinflammation, and cognitive decline (Aravindraja et al. 2022). Large-scale longitudinal data reveal that older adults with periodontal disease and oral infections are more likely to develop AD, with *Pg*-specific antibodies associated with both AD diagnosis and AD-related mortality (Merchant et al. 2024). Collectively, these findings support a strong association between periodontitis and AD, yet the mechanisms by which periodontitis contributes to AD development remain poorly understood.

Microglia, the resident immune cells of the CNS, play key roles in maintaining brain homeostasis by responding to injury or infection and by removing harmful proteins such as Aβ through phagocytosis and enzymatic degradation (Heneka et al. 2015). Under chronic stress, aging, or persistent inflammation, microglia lose their homeostatic state and become dysfunctional. This shift is marked by impaired phagocytosis, excessive production of inflammatory mediators, and increased oxidative stress (Mirarchi et al. 2024). Microglia dysfunction is a major contributor to AD pathology. We have previously shown that *Pg* can activate microglia and trigger robust inflammatory responses (Hao et al. 2022). However, the specific mechanisms through which *Pg* disrupts microglial function remain largely unknown.

Lipid droplets (LD) are dynamic, lipid-rich organelles consisting of a hydrophobic core of neutral lipids surrounded by a phospholipid monolayer and regulatory proteins (Jin et al. 2023). Once viewed as inert lipid storage sites, LD are now recognized as key regulators of cellular stress, metabolism, and inflammation (Verma et al. 2025). Under physiological conditions, LD formation is thought to exert protective effects by sequestering excess or toxic lipids and buffering cells against metabolic and oxidative stress (Cortini et al. 2025). However, when LD biogenesis becomes dysregulated, their accumulation can shift from adaptive to pathological (Yu et al. 2025). Aberrant or ectopic LD accumulation in non-adipose cells and tissues has been documented in a range of chronic inflammatory and metabolic disorders, including atherosclerosis, diabetes, obesity, rheumatoid arthritis, and AD (Chait and den Hartigh 2020), all of which have been linked with periodontitis (Hajishengallis 2022). In myeloid cells, abnormal LD accumulation has been linked with inflammation and cellular dysfunction (Zhao et al. 2024). This connection has gained particular attention in the CNS, where recent work has identified a distinct population of LD-accumulating microglia that emerges with aging (Marschallinger et al. 2020b). These LD-accumulating microglia exhibit a dysfunctional and proinflammatory state in the aging brain and contribute to neurodegenerative processes. *Pg* is known to disseminate systemically to influence neuroinflammatory pathways (Hao et al. 2022), yet it is not known if *Pg* affects LD accumulation in microglia and the functional consequences.

In the present study, we demonstrate that the *Pg* infection increases LD accumulation in microglia both in vitro and in vivo in an AD mouse model. These LD-accumulated microglia exhibit increased ROS production and impaired phagocytic capacity toward Ab. Inhibiting *Pg*- induced LD accumulation decreased ROS production and restored phagocytosis deficits of microglia. Our findings suggest that, in the context of periodontitis, systemic dissemination of periodontal pathogens may induce LD accumulation in microglia, which could establish a self-reinforcing cycle of microglia dysfunction that accelerates AD progression.

## Materials & Methods

Detailed descriptions of materials and methods are provided in the supplementary appendix.

### Mice

APP^NL-G-F/NL-G-F^ knock-in (*App* KI) mice were generated and maintained as previously described (Saito et al. 2014). All animal procedures were approved by the UAB Institutional Animal Care and Use Committee (IACUC) and were approved under protocol number 22766.

## Results

### *Pg* invades microglial cells and induces LD accumulation

Invasion to host cells and tissues is a critical virulence factor for *Pg*, enabling deeper tissue destruction, systemic spread, and immune evasion (Olsen and Progulske-Fox 2015). *Pg* has been shown to invade multiple cells, including epithelial cells, endothelial cells, and macrophages (Li et al. 2008). To evaluate *Pg* microglial invasion, CFSE-labeled *Pg* uptake by BV2 cells was analyzed by confocal microscopy. Distinct uptake of *Pg* in the cytoplasm of BV2 cells were observed 3 h following *Pg* treatment **(Figure 1A)**. The presence of *Pg* in BV2 cells was further validated by the presence of a significantly higher level of 16S rRNA expression in *Pg*-infected BV2 cells as compared with control cells **(Figure 1B)**.

**Figure 1.**
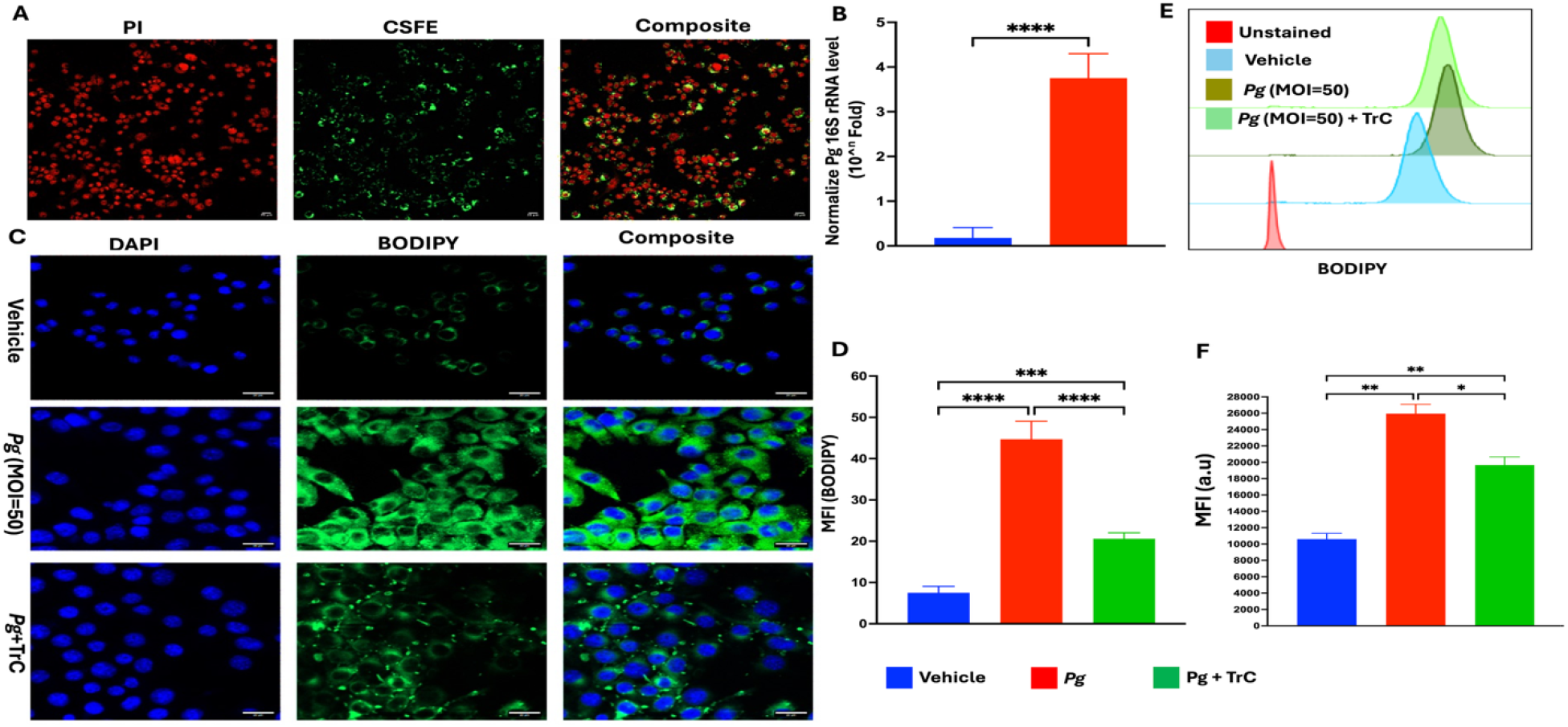
*Pg* invades BV2 microglial cells and induces LD accumulation. **(A)** Representative confocal fluorescent images of *Pg* invasion of BV2 cells. BV2 microglial cells were infected with CFSE-labeled *Pg* (green) at an MOI of 50 for 3 h, followed by PI staining (red). **(B)** The invasion of *Pg* in BV2 cells was confirmed by measuring *Pg* 16s rRNA level in BV2 microglial cells using RT-qPCR. **(C)** Representative confocal microscopy images of LD accumulation in BV2 cells. BV2 microglial cells were treated with TrC (1.0 µm) or PBS for 3 h, followed by *Pg* (MOI = 50) treatment for another 3h. Cells were primarily stained with BODIPY 493/503 (green) and counterstained with DAPI (blue); scale bar 20µm. **(D)** Quantification of the mean fluorescence intensity (MFI) of BODIPY 493/503 signal in a region of interest. **(E-F)** Representative flow cytometry histograms of BODIPY analysis and quantification of the MFI of BODIPY staining in BV2 cells. Data are presented as the mean ± SEM (n =3). *, *P* < 0.05, ** *P*< 0.01, *** *P* < 0.001, **** *P* < 0.0001 by one-way ANOVA.

To investigate time-dependent effect of *Pg*-induced LD formation, BV2 cells were infected with *Pg* (MOI = 50) for 0-24 h. Significantly increased LD accumulation was seen as early as 3 h post-infection **(Appendix figure 1)**. No further increase in LD number and intensity was noted at later time points. To investigate whether there is a dose-dependent effect, BV2 cells were infected with *Pg* at different MOIs for 3 h, and no significant differences in LD accumulation were seen at different doses of *Pg* tested **(Appendix figure 2)**.

To further validate LD accumulation, BV2 cells were pre-treated with TrC, an enzyme that is essential for the synthesis of triglycerides and cholesterol esters required for LD formation (Namatame et al. 1999). Results from both confocal microscopy and flow cytometry analysis showed that *Pg*-infection significantly increased LD accumulation in BV2 cells, while pre-treating cells with TrC significantly inhibited *Pg*-induced LD accumulation **(Figure 1C-F)**. These results demonstrate that *Pg* can invade microglial cells and induce LD accumulation.

### Inhibiting LD accumulation reduces *Pg*-induced oxidative stress in microglial cells

Emerging evidence indicates that brain tissue in patients with AD is subjected to oxidative stress, characterized by an imbalance between ROS production and impaired antioxidant defense mechanisms (Alkhalifa et al. 2025). Increased ROS production in microglia has been linked to microglial dysfunction and AD progression (Li et al. 2022). Here, we investigated the effect of *Pg* infection on ROS production and the role of LD in regulating the effect. Confocal microscopy revealed that *Pg* stimulation markedly increased ROS production compared to the controls **(Figure 2A-B)**. However, pre-treatment with TrC significantly reduced ROS levels, which is comparable to that observed with the ROS inhibitor AD4. Consistently, flow cytometry analysis showed that *Pg* stimulation significantly elevated ROS production, while TrC pre-treatment significantly attenuated this response **(Figure 2C-D)**.

**Figure 2.**
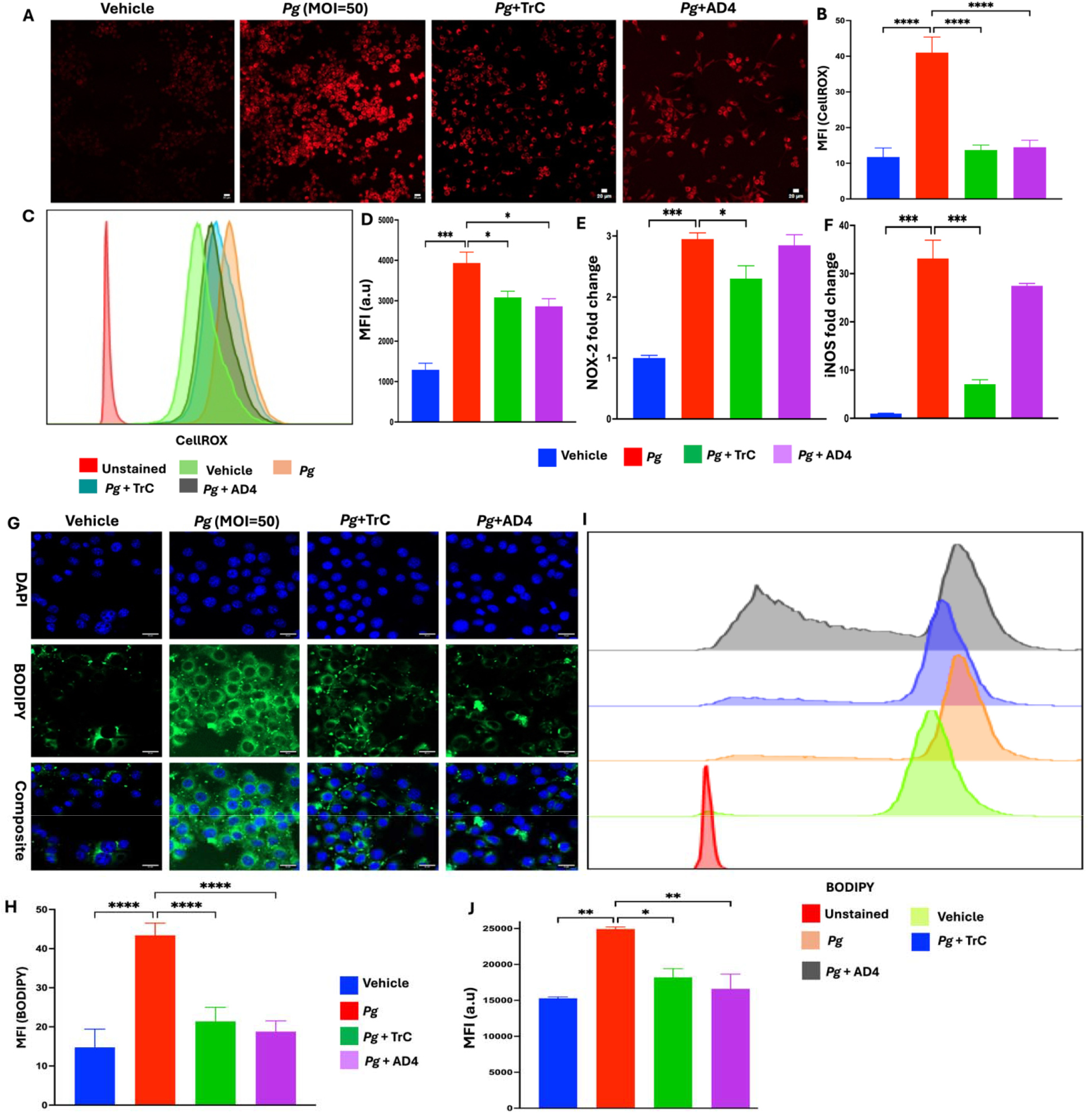
Bidirectional effects of *Pg*-induced LD accumulation and oxidative stress in BV2 microglial cells. BV2 microglial cells were treated with TrC (1.0 µm) and N-Acetylcysteine amide AD4 (500 µm) or vehicle for 3 h and then treated with *Pg* (MOI = 50) for another 3 h. **(A)** Representative confocal microscopy images showing CellROX^+^ in *Pg* -treated BV2 microglial cells. scale bar 20µm. **(B)** Quantification of the MFI of CellROX™ Deep Red signal in a region of interest. **(C)** Representative flow cytometry histograms of CellROX™ Deep Red staining of BV2 cells. **(D)** Quantification of the MFI of CellROX™ Deep Red staining. **(E-F)** NOX-2 and iNOS mRNA levels in BV2 cells expressed as fold-change over vehicle. **(G)** Representative confocal microscope images of BV2 cells stained with BODIPY 493/503 (Green) and DAPI (Blue). Scale bar 20µm. **(H)** Quantification of BODIPY 493/503 signal from confocal microscopy images. **(I-J)** Representative flow cytometry histograms of BODIPY 493/503 staining and quantification of BODIPY 493/503 staining in BV2 cells. Data are presented as the mean ± SEM (n = 3). *, *P* < 0.05, *** *P* < 0.001, **** *P* < 0.0001 by one-way ANOVA.

We next examined the expression of iNOS and NOX-2 genes, which are crucial producers of ROS (Nocella et al. 2023). *Pg*-infection significantly increased iNOS and NOX-2 mRNA expression in BV2 cells **(Figure 2E-F)**. Furthermore, pre-treatment with TrC markedly reduced iNOS and NOX-2 gene expression, while no significant inhibition of iNOS and NOX-2 gene expression was observed in BV2 cells pre-treated with AD4. It has been shown that AD4 removes ROS that are already produced via boosting intracellular glutathione level, but it does not necessarily suppress the signaling pathways that drive iNOS/NOX-2 gene expression (Amer et al. 2008). Taken together, our results indicate that *Pg*-induced LD accumulation contributes to oxidative stress in microglial cells via regulation of iNOS and NOX-2 genes.

### *Pg*-induced oxidative stress regulates LD accumulation in microglial cells

Recent studies have shown that oxidative stress promotes the biogenesis of LD, which protects vulnerable lipids such as unsaturated fatty acids by rerouting them into the triglyceride core (Cortini et al. 2025). This sequestration prevents ROS-induced peroxidation, thereby preserving lipid homeostasis and underscoring the regulatory role of oxidative stress on LD-production (Lange et al. 2025). To investigate whether *Pg*-induced oxidative stress reciprocally regulates LD, the effect of AD4 on *Pg*-induced LD accumulation was examined. AD4 pre-treatment markedly reduced LD accumulation in BV2 cells compared to the *Pg*-infected group **(Figure 2G-H)**. This effect was comparable to that observed with the LD inhibitor TrC. Flow cytometry analysis confirmed that pre-treatment with TrC and AD4 significantly reduced LD accumulation in BV2 cells **(Figure 2I-J)**. These results indicate that *Pg*-infection induces both oxidative stress and LD accumulation in microglial cells, and that a reciprocal regulatory relationship exists between oxidative stress and LD formation following *Pg* stimulation.

### *Pg*-induced LD accumulation in microglial cells impairs their ability to phagocytose Aβ peptides

One of the primary functions of microglia is to clear cellular debris, including Aβ peptides, whose buildup contributes to cognitive decline and AD (Brown et al. 2026). To investigate the effect of *Pg* on microglial phagocytosis of Ab peptides and the role of LD in regulating the effect, BV2 cells were pre-treated with vehicle or TrC, followed by *Pg-*infection, and their ability to phagocytose Hilyte Fluor 488-labeled Aβ was analyzed. CytoD, an inhibitor of actin polymerization, was used as a positive control for phagocytosis inhibition (Xie et al. 2025). Confocal microscopy showed that vehicle-treated BV2 cells exhibited the highest level of Aβ internalization, whereas CytoD significantly reduced Aβ uptake as compared to the other groups **(Figure 3A-B)**. *Pg* infection significantly reduced Aβ uptake in BV2 cells, while TrC restored the uptake. Flow cytometry further confirmed that the vehicle-treated cells had the maximal rate of Aβ internalization, while *Pg* infection significantly reduced the Aβ uptake, and TrC restored the Aβ uptake **(Figure 3C-D)**. Taken together, these results demonstrate that *Pg* infection impairs microglia phagocytosis of Aβ, and that inhibition of LD can restore the ability of microglial cells to phagocytose Aβ.

**Figure 3.**
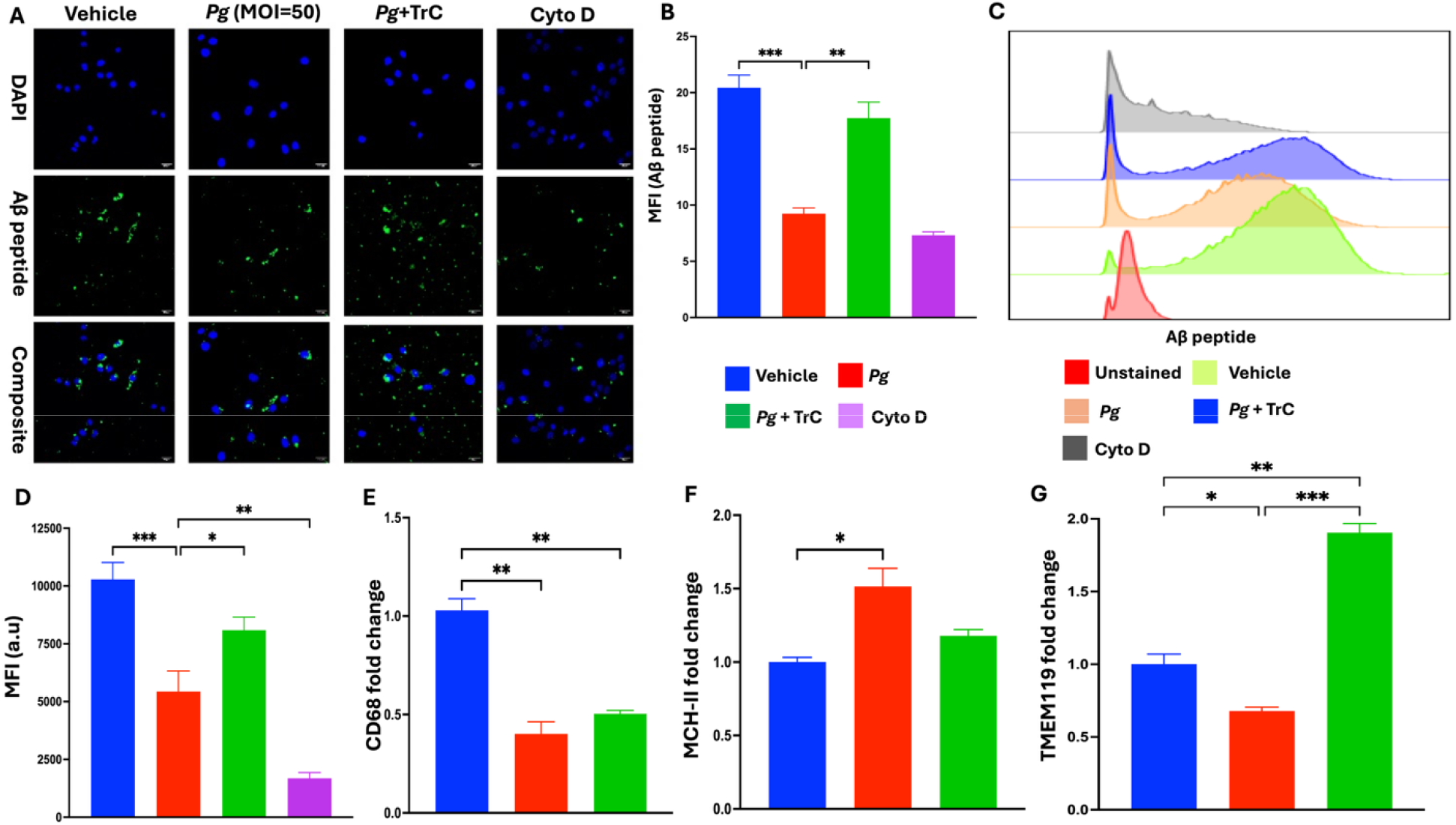
*Pg*-induced LD impairs microglia phagocytosis of Aβ. BV2 microglia were pre-treated with TrC and Cyto D for 3 h and then infected with *Pg* (MOI = 50) for an additional 3 h, followed by incubation with Aß (1-42) Hilyte flour 488 for 2 h. **(A)** Representative confocal microscopy images of BV2 microglial cells stained with Aß (1-42) Hilyte flour 488 (Green) and DAPI (Blue). Scale bar 20µm. **(B)** Quantification of Aß (1-42) Hilyte flour 488 signals from confocal microscope images. **(C-D)** Representative flow cytometry histograms and quantification of Aß (1-42) Hilyte flour 488 in BV2 cells. **(E-G)** The relative mRNA levels of CD68, MCH-II, and TMEM119 in BV2 cells. Data are presented as the mean ± SEM (n = 3). *, *P* < 0.05, ** *P* < 0.01, *** *P* < 0.001 by one-way ANOVA.

Since microglial phagocytosis ability is directly linked to their activation state (Valiukas et al. 2025), we next evaluated the activation state of BV2 cells following *Pg*-infection. *Pg* infection significantly downregulated the gene expression of CD68, a key marker for activated and phagocytic microglia, as compared to control cells **(Figure 3E**). Interestingly, the expression of MHC-II, another microglial activation marker, was significantly up-regulated following *Pg* stimulation, while the expression of TMEM119, a microglial resting marker, was significantly down-regulated **(Figure 3F-G)**. No difference in CD68 gene expression was seen following TrC treatment compared to *Pg*-treated cells **(Figure 3E)**. These results suggest that *Pg*-infection causes microglia dysfunction, characterized by increased activation and reduced phagocytosis, at least partially due to *Pg*-induced LD accumulation.

### *Pg*-infection enhances LD accumulation and impairs the functionality of microglia in *App* KI mice

We next assessed if *Pg*-infection could induce LD accumulation in microglia in vivo, and if these affect oxidative stress and microglial phagocytosis of Ab. *App* KI mice (Pang et al. 2022) were infected with *Pg* or vehicle via retro-orbital injection, and brain tissues were collected 6 h post-infection for further analysis **(Appendix figure 3)**. For confocal microscopy analysis, brain sections were co-stained with IBA1 (microglia), LipidSpot (LD), and Methoxy-X04 (Aβ) to examine the spatial distribution of microglia, LD, and Ab plaques. Since the hippocampus is significantly affected by Ab pathology in AD patients and mouse models (Vyas et al. 2020), we focused our analysis on this region. An age-dependent increase in the number of hypertrophic microglia with enlarged cell bodies, along with an increased number and size of LD, were observed in controls. *Pg* infection further increased the number of LD and the LD-laden microglia at 3 months of age. Importantly, Aβ plaques were predominantly adjacent to LD-laden microglia, and *Pg* infection significantly increased the number of LD-loaded microglia adjacent to Aβ plaques **(Appendix figure 4A-D)**. Similar findings were seen in mice at 6 and 12 months of age **(Appendix figure 4E-L)**. The number of hypertrophic microglia with enlarged cell bodies was also observed in *Pg*-infected *App* KI mice compared with control *App* KI mice **(Figure 4A)**. These results indicate that *Pg* infection increases LD in microglia *in vivo*.

**Figure 4.**
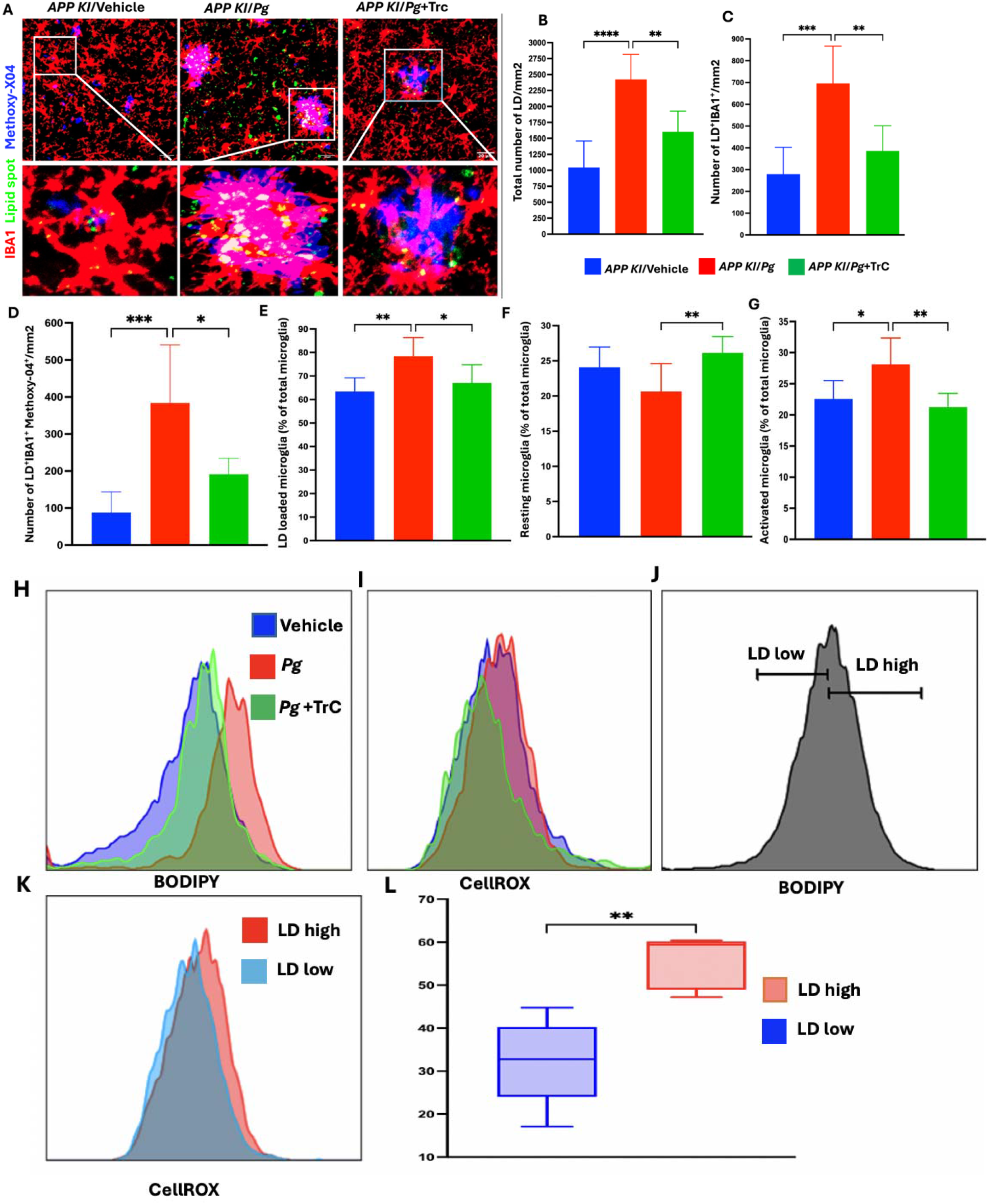
*Pg*-induced LD accumulation impairs phagocytosis and enhancing ROS production in microglia in *App* KI mice. **(A)** Representative confocal image of microglia (IBA1; red), LD (LipidSpot™ 488; green), and Ab peptides (Methoxy-X04; blue) from hippocampus of *App KI* mice with *Pg* or without *Pg* infection. Scale bar: 20µm. **(B-D)** Quantification of total LD, LD-loaded microglia, and LD-loaded microglia at Aß positive area on confocal microscopy images. **(E-G)** Quantification of flow cytometry analysis of total LD-loaded microglia, resting microglia, and activated microglia. **(H-I)** Represented flow histogram of LD- and CellROX-accumulating microglia from the brains of *App KI* mice. **(J)** The flow gating scheme for LD-high and LD-low microglia from the brains of *App KI* mice. **(K-L)** Representative histogram and quantification of CellROX fluorescence in LD high and LD low microglia from the brains of *App KI* mice. Data are presented as the mean ± SEM (n =6). *, *P* < 0.05, ** *P* < 0.01, *** *P* < 0.001 by one-way ANOVA and Welch’s t test.

To investigate how LD accumulation regulates microglial activation and phagocytosis, *App KI* mice were pre-treated with TrC before *Pg* infection. Confocal microscopy showed that TrC significantly reduced the number of LD-loaded microglia and their proximity to Aβ plaques **(Figure 4A-D)**. Similarly, flow cytometry analysis showed that Pg infection increased the percentage of LD-loaded microglia and activated microglia without affecting the total number of microglia **(Figure 4E-G)**. TrC treatment significantly decreased the percentage of LD-loaded microglia and activated microglia. Taken together, these results indicate that *Pg* infection increases LD accumulation in microglia in AD mice and that *Pg*-induced LD accumulation impairs microglial phagocytosis of Ab in *vivo*.

To investigate how LD accumulation influences microglial oxidative stress in AD mice following *Pg* infection, we analyzed ROS intensity in microglia from *App KI* mice. *Pg* infection elevated LD and intracellular ROS levels **(Figure 4H-I)**. Notably, within the *Pg*-infected group, microglia with high LD content exhibited significantly higher ROS intensity than those with low LD content **(Figure 4J-L)**. Furthermore, inhibition of LD by TrC treatment reduced ROS intensity. Collectively, these findings suggest that *Pg* infection promotes LD accumulation in microglia in AD mice, which contributes to the oxidative stress of microglia.

## Discussions

Microglia are the primary innate immune cells in the brain (El Mesaoudi et al. 2026), and growing evidence indicates that LD accumulation in microglia represents a key pathological feature of AD, linking metabolic dysfunction, impaired immune responses, and neurodegeneration (Wu et al. 2025). For many years, the precise role of microglia in brain pathology has been subject to debate (Chakrabarty et al. 2015). Currently, microglia are recognized as immune sentinels and active sculptors of the brain microenvironment, influencing synaptic remodeling, neuronal survival, and tissue homeostasis (Ye et al. 2025). In AD, microglia undergo profound metabolic and functional changes, and the emergence of LD-rich microglia has recently been identified as a hallmark of aging and neurodegeneration (Xing et al. 2025). While LD formation may initially serve adaptive roles, excessive LD accumulation has been shown to impair microglial functions (Li et al. 2025).

Increasing evidence suggests that periodontitis is associated with a range of systemic disorders, including AD (Chen et al. 2026). Although the mechanisms responsible for this association remain largely unknown, the key periodontal pathogen *Pg* has been detected in AD brains and linked to elevated Aβ and tau pathology (Dominy et al. 2019). However, there is no information if *Pg* exerts any influence on LD accumulation in microglia and the metabolic consequences of this effect. Our study addresses this gap by demonstrating that *Pg* infection is a potent inducer of LD accumulation in microglia, both *in vitro* in BV2 cells and *in vivo* in *App* KI mice. To our knowledge, this is the first report demonstrating that *Pg* infection enhances LD accumulation in microglia within an AD-relevant animal model. These findings align with literature showing that *Pg* infection can induce LD formation in macrophages (Rho et al. 2021) and that chronic periodontitis contributes to foam cell formation in peripheral immune cells (Afzoon et al. 2023).

One potential mechanism linking LD accumulation to neurodegeneration is lipid peroxidation (Sung et al. 2025). When exposed to ROS, LD can undergo peroxidation, generating cytotoxic lipid species that exacerbate cellular stress and contribute to neuronal injury (Liu et al. 2015). Enhanced ROS generation is a hallmark of LD-rich microglia and has also been observed in LD-loaded peripheral immune cells (Marschallinger et al. 2020b), suggesting a conserved stress-associated metabolic phenotype across immune compartments. However, whether ROS drives LD accumulation or vice versa remains debated. Some studies propose that oxidative stress promotes LD biogenesis as a protective hub for damaged lipids (Jarc and Petan 2019), whereas others argue that excessive LD buildup can itself enhance ROS generation through impaired mitochondrial function or increased substrate availability for lipid peroxidation (Sung et al. 2025). Our results suggest a bidirectional relationship between LD accumulation and ROS generation in the presence of *Pg* infection: inhibition of LD formation with TrC markedly suppressed ROS generation in BV2 cells and in microglia in AD mice, indicating that LD contributes to *Pg*-induced oxidative stress; conversely, pharmacologic ROS inhibition reduced LD accumulation, suggesting that early ROS production may initiate LD formation. This bidirectional relationship may potentially create a feed-forward loop that accelerates neurodegenerative processes.

Microglial phagocytosis is essential for clearing cellular debris, protein aggregates, and dysfunctional synapses that accumulate with aging (Brown et al. 2026). When this clearance function declines, toxic species, such as Aβ build up, promoting neuronal damage in AD (Dias and Socodato 2025). Impaired microglial phagocytosis is therefore recognized as a key contributor to AD progression. Recent studies have identified phagosome maturation as the top-regulated pathway in LD-accumulating microglia, and these cells exhibit marked deficits in zymosan particle uptake (Marschallinger et al. 2020a). In our work, we found that *Pg* infection similarly disrupts microglial function. We also observed an age-dependent increase in LD-loaded, plaque-proximal microglia in *App KI* mice. This aligns with recent reports showing that LD-laden plaque-proximal microglia are present in both human AD brain and the Ab-rich 5xFAD mouse model (Prakash et al. 2025). Notably, *Pg* infection further amplified the number of LD-rich microglia in proximity to Aβ plaques. Together, these findings support a model in which *Pg* infection exacerbates microglial LD accumulation, thereby impairing phagocytosis and potentially accelerating AD-related pathology.

Microglial activation is another key factor linking LD-accumulation with impaired microglial function (Konishi et al. 2020). We have shown previously that *Pg* infection induces microglia activation in vitro and exacerbates microglia activation in vivo in AD mice (Hao et al. 2022); however, the functionality of these activated microglia needs further study. Data from our present study demonstrate that *Pg* infection upregulates microglial activation markers in parallel with LD accumulation and phagocytic impairment, suggesting that metabolic and functional activation are intertwined. Importantly, TrC treatment not only reduced LD-accumulation but also restored microglia homeostasis, highlighting the potential of targeting lipid metabolism to improve microglial function. Indeed, recent studies have suggested that restoring microglial homeostasis may be a promising therapeutic strategy for slowing AD progression (Liu et al. 2025). Our ongoing studies are exploring the signaling mechanisms involved in *Pg*-induced LD accumulation in microglia.

Taken together, our data support a model in which periodontal infection drives LD accumulation in microglia, and this metabolic alteration exacerbates microglia dysfunction via a self-reinforcing cycle of excessive oxidative stress and impaired phagocytic function that may accelerate AD progression **(Figure 5)**. While epidemiological studies have long suggested an association between periodontitis and cognitive decline, our findings offer a mechanistic explanation rooted in microglial metabolism. Our work strengthens the emerging oral-brain axis and highlights the importance of maintaining oral health in improving the progression of neurodegenerative diseases, including AD.

**Figure 5.**
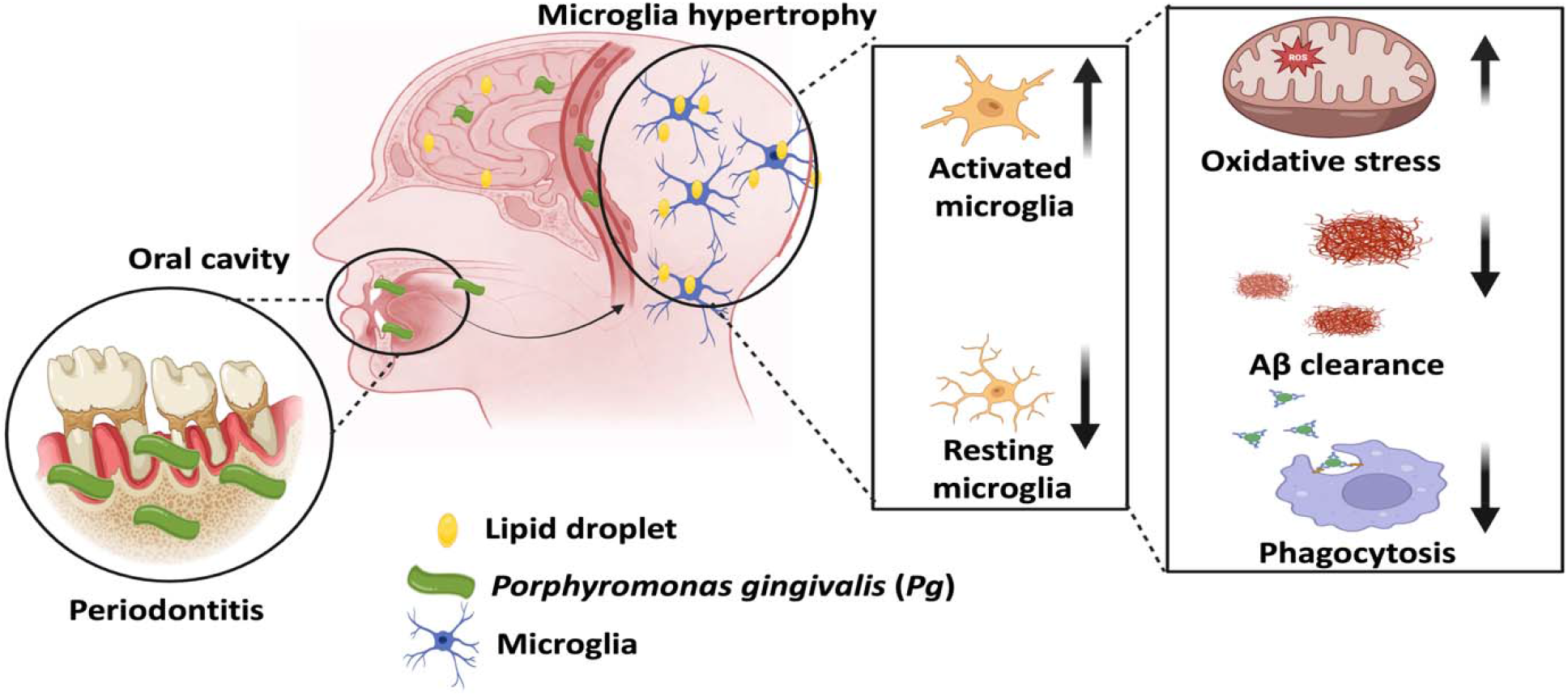
Schematic illustration of *Pg*-induced LD accumulation in microglia in accelerating AD progression. In the context of periodontitis, systemic dissemination of periodontal pathogens such as *Pg* may induce LD accumulation in microglia, which could establish a self-reinforcing cycle of microglia dysfunction by elevating ROS production and impair Aβ clearance, thus accelerating AD progression.

## AUTHOR CONTRIBUTIONS

Muhammad Shahid Riaz Rajoka and Ping Zhang contributed to conception and design, data acquisition, analysis, and interpretation, drafted and critically revised the manuscript. Kristina Nicole Valladares, Chloe La Prairie, and Wei Li contributed to data acquisition, analysis, and interpretation, and critically revised the manuscript. Peter King, Jannet Katz, and Suzanne M. Michalak contributed to conception and design, data analysis, and interpretation, and critically revised the manuscript. All authors gave their final approval and agreed to be accountable for all aspects of the work.

## ACKNOWLEDGMENTS

We thank Greg Harber for his technical assistance. We also acknowledge the UAB Comprehensive Flow Cytometry Core, the UAB Comprehensive Neuroscience Center, and the UAB Civitan International Research Center for their technical support during data acquisition.

## Declaration of Conflicting Interests

The authors declared no potential conflicts of interest with respect to the research, authorship, and/or publication of this article.

## Funding

This study was supported by grants from National Institute of Dental and Craniofacial Research (NIDCR) (R01 DE026465 to P.Z) and National Institute of Aging (NIA) (R01 AG078708 to P.Z), and a pilot grant from UAB Global Center for Craniofacial, Oral and Dental Disorders (GC-CODED to P. Z.). M.S.R.R., K. N. V., and C. L.P are supported by NIDCR Dental Academic Research Training grant (R90 DE023056). P.H.K is supported by the U.S. Department of Veterans Affairs grant (BX006231). UAB Comprehensive Flow Cytometry Core is supported by the National Institutes of Health grants P30AI27667 and P30AR048311.

## DATA AVAILABILITY STATEMENT

All data supporting the findings of this study are included in the article and its Appendix.

## Notes

### Competing Interest Statement

The authors have declared no competing interest.

## References

Afzoon S, Amiri MA, Mohebbi M, Hamedani S, Farshidfar N. 2023. A systematic review of the impact of porphyromonas gingivalis on foam cell formation: Implications for the role of periodontitis in atherosclerosis. BMC Oral Health. 23(1):481.

Alkhalifa AE, Alkhalifa O, Durdanovic I, Ibrahim DR, Maragkou S. 2025. Oxidative stress and mitochondrial dysfunction in alzheimer’s disease: Insights into pathophysiology and treatment. Journal of Dementia and Alzheimer's Disease. 2(2):17.

Amer J, Atlas D, Fibach E. 2008. N-acetylcysteine amide (ad4) attenuates oxidative stress in beta-thalassemia blood cells. Biochimica et Biophysica Acta (BBA) - General Subjects. 1780(2):249–255.

Aravindraja C, Sakthivel R, Liu X, Goodwin M, Veena P, Godovikova V, Fenno JC, Levites Y, Golde TE, Kesavalu L. 2022. Intracerebral but not peripheral infection of live porphyromonas gingivalis exacerbates alzheimer’s disease like amyloid pathology in app-tgcrnd8 mice. International Journal of Molecular Sciences. 23(6):3328.

Brown GC, St George-Hyslop P, Paolicelli RC, Lemke G. 2026. Microglial phagocytosis in alzheimer disease. Nature Reviews Neurology. 22(1):54–69.

Cavaco MA, Jang SR, Olsen C, Bodnar C, Ferko N. 2025. Global societal burden of alzheimer’s disease by severity: A targeted literature review. Neurol Ther. 14(5):1797–1826.

Chait A, den Hartigh LJ. 2020. Adipose tissue distribution, inflammation and its metabolic consequences, including diabetes and cardiovascular disease. Front Cardiovasc Med. 7:22.

Chakrabarty P, Li A, Ceballos-Diaz C, Eddy James A, Funk Cory C, Moore B, DiNunno N, Rosario Awilda M, Cruz Pedro E, Verbeeck C et al. 2015. Il-10 alters immunoproteostasis in app mice, increasing plaque burden and worsening cognitive behavior. Neuron. 85(3):519–533.

Chen C, Chen Q, Zou H, Zhao C, Wang X. 2026. Research trends in periodontitis and alzheimer’s disease: A bibliometric analysis based on web of science and scopus. International Dental Journal. 76(1):109327.

Cortini M, Ilieva E, Massari S, Bettini G, Avnet S, Baldini N. 2025. Uncovering the protective role of lipid droplet accumulation against acid-induced oxidative stress and cell death in osteosarcoma. Biochimica et Biophysica Acta (BBA) - Molecular Basis of Disease. 1871(2):167576.

Dias D, Socodato R. 2025. Beyond amyloid and tau: The critical role of microglia in alzheimer’s disease therapeutics. Biomedicines. 13(2):279.

Dominy SS, Lynch C, Ermini F, Benedyk M, Marczyk A, Konradi A, Nguyen M, Haditsch U, Raha D, Griffin C et al. 2019. porphyromonas gingivalis in alzheimer’s disease brains: Evidence for disease causation and treatment with small-molecule inhibitors. Science Advances. 5(1):eaau3333.

El Mesaoudi A, Kassoussi A, Zahaf A, Ayadi M, Naglieri S, Marie C, Razavi F, Bobé P, Martinovic J, Parras C et al. 2026. Smoothened-mediated signaling contributes to immune and non-immune functions of microglia. Brain, Behavior, and Immunity. 132:106226.

Hajishengallis G. 2022. Interconnection of periodontal disease and comorbidities: Evidence, mechanisms, and implications. Periodontol 2000. 89(1):9–18.

Hao X, Li Z, Li W, Katz J, Michalek SM, Barnum SR, Pozzo-Miller L, Saito T, Saido TC, Wang Q et al. 2022. Periodontal infection aggravates c1q-mediated microglial activation and synapse pruning in alzheimer’s mice. Frontiers in Immunology. Volume 13 -2022.

Heneka MT, Golenbock DT, Latz E. 2015. Innate immunity in alzheimer’s disease. Nature Immunology. 16(3):229–236.

Jarc E, Petan T. 2019. Lipid droplets and the management of cellular stress. Yale J Biol Med. 92(3):435–452.

Jin Y, Tan Y, Wu J, Ren Z. 2023. Lipid droplets: A cellular organelle vital in cancer cells. Cell Death Discovery. 9(1):254.

Klyucherev TO, Olszewski P, Shalimova AA, Chubarev VN, Tarasov VV, Attwood MM, Syvänen S, Schiöth HB. 2022. Advances in the development of new biomarkers for alzheimer’s disease. Transl Neurodegener. 11(1):25.

Knopman DS, Amieva H, Petersen RC, Chételat G, Holtzman DM, Hyman BT, Nixon RA, Jones DT. 2021. Alzheimer disease. Nature Reviews Disease Primers. 7(1):33.

Konishi H, Okamoto T, Hara Y, Komine O, Tamada H, Maeda M, Osako F, Kobayashi M, Nishiyama A, Kataoka Y et al. 2020. Astrocytic phagocytosis is a compensatory mechanism for microglial dysfunction. The EMBO Journal. 39(22):e104464.

Lamont RJ, Hajishengallis G. 2015. Polymicrobial synergy and dysbiosis in inflammatory disease. Trends in Molecular Medicine. 21(3):172–183.

Lange M, Wölk M, Doubravsky CE, Hendricks JM, Kato S, Otoki Y, Styler B, Nakagawa K, Fedorova M, Olzmann JA. 2025. Fsp1-mediated lipid droplet quality control prevents neutral lipid peroxidation and ferroptosis. bioRxiv.

Laugisch O, Johnen A, Maldonado A, Ehmke B, Bürgin W, Olsen I, Potempa J, Sculean A, Duning T, Eick S. 2018. Periodontal pathogens and associated intrathecal antibodies in early stages of alzheimer’s disease. Journal of Alzheimer’s Disease. 66(1):105–114.

Li L, Michel R, Cohen J, DeCarlo A, Kozarov E. 2008. Intracellular survival and vascular cell-to-cell transmission of porphyromonas gingivalis. BMC Microbiology. 8(1):26.

Li Y, Xia X, Wang Y, Zheng JC. 2022. Mitochondrial dysfunction in microglia: A novel perspective for pathogenesis of alzheimer’s disease. Journal of Neuroinflammation. 19(1):248.

Li Y, Zhao Q, Wang Y, Du W, Yang R, Wu J, Li Y. 2025. Lipid droplet accumulation in microglia and their potential roles. Lipids Health Dis. 24(1):215.

Liu J, Wang Z, Liang W, Zhang Z, Deng Y, Chen X, Hou Z, Xie Y, Wang Q, Li Y et al. 2025. Microglial tmem119 binds to amyloid-β to promote its clearance in an aβ-depositing mouse model of alzheimer’s disease. Immunity. 58(7):1830-1846.e1837.

Liu L, Zhang K, Sandoval H, Yamamoto S, Jaiswal M, Sanz E, Li Z, Hui J, Graham Brett H, Quintana A et al. 2015. Glial lipid droplets and ros induced by mitochondrial defects promote neurodegeneration. Cell. 160(1):177–190.

Marschallinger J, Iram T, Zardeneta M, Lee SE, Lehallier B, Haney MS, Pluvinage JV, Mathur V, Hahn O, Morgens DW et al. 2020a. Lipid-droplet-accumulating microglia represent a dysfunctional and proinflammatory state in the aging brain. Nat Neurosci. 23(2):194–208.

Marschallinger J, Iram T, Zardeneta M, Lee SE, Lehallier B, Haney MS, Pluvinage JV, Mathur V, Hahn O, Morgens DW et al. 2020b. Lipid-droplet-accumulating microglia represent a dysfunctional and proinflammatory state in the aging brain. Nature Neuroscience. 23(2):194–208.

Merchant AT, Zhao L, Bawa EM, Yi F, Vidanapathirana NP, Lohman M, Zhang J. 2024. Association between clusters of antibodies against periodontal microorganisms and alzheimer disease mortality: Evidence from a nationally representative survey in the USA. Journal of Periodontology. 95(1):84–90.

Mirarchi A, Albi E, Arcuri C. 2024. Microglia signatures: A cause or consequence of microglia-related brain disorders? International Journal of Molecular Sciences. 25(20):10951.

Namatame I, Tomoda H, Arai H, Inoue K, Omura S. 1999. Complete inhibition of mouse macrophage-derived foam cell formation by triacsin c. The Journal of Biochemistry. 125(2):319–327.

Nocella C, D’Amico A, Cammisotto V, Bartimoccia S, Castellani V, Loffredo L, Marini L, Ferrara G, Testa M, Motta G et al. 2023. Structure, activation, and regulation of nox2: At the crossroad between the innate immunity and oxidative stress-mediated pathologies. Antioxidants. 12(2):429.

Olsen I, Progulske-Fox A. 2015. Invasion of porphyromonas gingivalis strains into vascular cells and tissue. Journal of Oral Microbiology. 7(1):28788.

Pang K, Jiang R, Zhang W, Yang Z, Li L-L, Shimozawa M, Tambaro S, Mayer J, Zhang B, Li M et al. 2022. An app knock-in rat model for alzheimer’s disease exhibiting aβ and tau pathologies, neuronal death and cognitive impairments. Cell Research. 32(2):157–175.

Passeri E, Elkhoury K, Morsink M, Broersen K, Linder M, Tamayol A, Malaplate C, Yen FT, Arab-Tehrany E. 2022. Alzheimer’s disease: Treatment strategies and their limitations. Int J Mol Sci. 23(22).

Prakash P, Manchanda P, Paouri E, Bisht K, Sharma K, Rajpoot J, Wendt V, Hossain A, Wijewardhane PR, Randolph CE et al. 2025. Amyloid-β induces lipid droplet-mediated microglial dysfunction via the enzyme dgat2 in alzheimer’s disease. Immunity. 58(6):1536-1552.e1538.

Rho JH, Kim HJ, Joo JY, Lee JY, Lee JH, Park HR. 2021. Periodontal pathogens promote foam cell formation by blocking lipid efflux. Journal of Dental Research. 100(12):1367–1377.

Saito T, Matsuba Y, Mihira N, Takano J, Nilsson P, Itohara S, Iwata N, Saido TC. 2014. Single app knock-in mouse models of alzheimer’s disease. Nature Neuroscience. 17(5):661–663.

Sung S, Kim H-J, Cha SJ, Nahm M, Kim SH, Kwon M-S. 2025. Microglial lipid droplets as therapeutic targets in age-related neurodegenerative diseases. npj Aging. 12(1):2.

Uhlmann RE, Rother C, Rasmussen J, Schelle J, Bergmann C, Ullrich Gavilanes EM, Fritschi SK, Buehler A, Baumann F, Skodras A et al. 2020. Acute targeting of pre-amyloid seeds in transgenic mice reduces alzheimer-like pathology later in life. Nat Neurosci. 23(12):1580–1588.

Valiukas Z, Tangalakis K, Apostolopoulos V, Feehan J. 2025. Microglial activation states and their implications for alzheimer’s disease. The Journal of Prevention of Alzheimer’s Disease. 12(1):100013.

Verma R, Sharma P, Sharma V, Singh TG. 2025. Modulating lipid droplet dynamics in neurodegeneration: An emerging area of molecular pharmacology. Molecular Biology Reports. 52(1):277.

Vyas Y, Montgomery JM, Cheyne JE. 2020. Hippocampal deficits in amyloid-β-related rodent models of alzheimer’s disease. Front Neurosci. 14:266.

Wu X, Miller JA, Lee BTK, Wang Y, Ruedl C. 2025. Reducing microglial lipid load enhances β amyloid phagocytosis in an alzheimer’s disease mouse model. Science Advances. 11(6):eadq6038.

Xie M, Huang X, Tang Q, Yu S, Zhang K, Lu X, Zhang Y, Wang J, Zhang L, Chen L. 2025. Porphyromonas gingivalis impairs microglial aβ clearance in a mouse model. J Dent Res. 104(4):408–418.

Xing J, McKenzie T, Hu J. 2025. Lipid-laden microglia: Characterization and roles in diseases. Cells. 14(16):1281.

Ye G, Wang Z, Chen P, Ye J, Li S, Chen M, Feng J, Wang H, Chen W. 2025. Serpina3n in neonatal microglia mediates its protective role for damaged adult microglia by alleviating extracellular matrix remodeling-induced tunneling nanotubes degradation in a cell model of traumatic brain injury. Neuroscience. 565:1–9.

Yu J, Liu Q, Zhang Y, Xu L, Chen X, He F, Zhang M, Yang H, Yu S, Liu X et al. 2025. Stress causes lipid droplet accumulation in chondrocytes by impairing microtubules. Osteoarthritis and Cartilage. 33(3):351–363.

Zhao R, Zhang L, Kamachi K, Gupta SK, Akiyama H, Nguyen J, Andreeff M, Ishizawa J, Lodi A. 2024. Isolation of lipid droplets from acute myeloid leukemia cells for lipidomic analysis. Blood. 144:7502.

