## Appendix materials for "*Porphyromonas gingivalis* promotes lipid droplet-mediated microglial dysfunction": JDR Supplementary appendix .docx

**Running Title:** *P. gingivalis* infection promotes microglial dysfunction

**Correspondence**

Ping Zhang, Department of Pediatric Dentistry, School of Dentistry, University of Alabama at Birmingham, 1919 7^th^ Avenue South, Birmingham, AL 35294, USA.

**Supplementary appendix detailed materials and methods**

**Bacterial culture**

*Porphyromonas gingivalis* (*Pg*) ATCC 33277 was grown on enriched trypticase soy agar plates containing 1% yeast extract, 5% defibrinated sheep blood, 5% hemin, and 1% menadione, at 37˚C in an anaerobic environment (10% H_2_, 5% CO_2_, and 85% N_2_) for 5-7 days, followed by subculturing overnight in trypsin soy broth supplemented with yeast (1%), hemin (5 µg/ml) and menadione (1 µg/ml) (Hao et al. 2022; Zhao et al. 2020).

**BV2 cell culture**

BV2 murine microglia cells (Amsbio; cat#AMS.EP-CL-0493) were grown in DMEM medium (Corning Incorporated, NY, USA) containing 10% FBS, 1% Penicillin/Streptomycin (P/S), and 1% L-glutamine at 37 °C in a 5% CO_2_ incubator (Yazdanpanah Moghadam et al. 2025).

**Mice**

APP^NL-G-F/NL-G-F^ knock-in (*App* KI) mice (Saito et al. 2014) that harbor the APP Swedish (KM670/671NL), Arctic (E693G), and Liberian (I716F) mutations were originally obtained from Dr. Takaomi Saido (RIKEN Brain Science Institute, Japan) under a material transfer agreement and had been backcrossed with C57BL/6 mice (Jackson Laboratories, Ellsworth, ME, USA) to obtain the *App* KI mice with a C57BL/6 background. All animals were housed in a controlled environment at the University of Alabama at Birmingham (UAB), and all experimental procedures were approved by the UAB Institutional Animal Care and Use Committee.

**In vitro detection of *Pg* invasion in BV2 cells**

To assess *Pg* invasion in BV2 cells, freshly prepared *Pg* were labeled with carboxyfluorescein succinimidyl ester (CFSE) (Dojindo Laboratories) by incubating the bacteria with 10 µM CFSE for 15 minutes (min) at room temperature (RT). The CFSE-labeled *Pg* were then washed and resuspended in PBS for subsequent experiments. BV2 cells were infected with CFSE-labeled *Pg* at an MOI of 50 for 3 hours (h), washed with PBS, and fixed with 4% paraformaldehyde (PFA) for 15 min. After fixation, BV2 cells were stained with propidium iodide (PI) for 30 min, washed with PBS, and mounted using Fluromount-G^TM^ mounting medium without DAPI (Thermo Fisher Scientific). The intracellular presence of *Pg* was visualized by confocal microscopy (Zeiss LSM-800).

In addition, *Pg* internalization was confirmed by the detection of *Pg* specific16S ribosomal (rRNA) by real-time reverse transcription quantitative PCR (RT-qPCR). Briefly, total RNA was extracted using Trizol reagent (Thermo Fisher Scientific), and 1 ug of total RNA was reverse transcribed using PrimeScript^TM^ RT Reagent Kit (Takara Bio). Real-time PCR was performed with TB Green^R^ Advantage^R^ qPCR premix (Takara Bio) in the ABI 7500 real-time PCR detection system (Applied Biosystems). The target gene expressions were normalized to GAPDH expression obtained from a parallel reaction. Primer sequences are listed in **Table S1.**

**Confocal microscopy analysis of LD accumulation in BV2 cells following *Pg* infection**

BV2 cells (5×10^4^ cells/well) were seeded on 8-well chamber slides (Thermo Fisher Scientific) and infected with *Pg* at various doses (MOI of 5, 10, 25, or 50) for different time periods (3, 6, 9, 12, or 24 h). To inhibit LD formulation, BV2 cells were pretreated with vehicle or 1.0 μM of triacsin C (TrC) (Cyman Chemical), an inhibitor of long fatty acyl-CoA synthetase, for 3 h, followed by *Pg* stimulation. For confocal microscopy analysis, BV2 cells were fixed with 4% PFA, washed three times with PBS, and incubated with BODIPY 493/503 (Thermo Fisher Scientific; 1:1000 from 1 mg/ml stock solution in DMSO) for 30 min at RT, protected from light. The cells were then washed three times with PBS and mounted using DAPI Fluoromount-G^R^ mounting medium (Southern Biotech). Images were acquired from randomly selected regions using a confocal microscope. The mean fluorescence intensity (MFI) of BODIPY 493/503 staining was quantified using ImageJ software (National Institute of Health).

**Flow cytometry analysis of LD accumulation in BV2 cells following *Pg* infection**

For flow cytometry analysis, BV2 cells (5×10^5^ cells/well) were seeded into 24-well plates, pretreated with vehicle or TrC for 3 h, and infected with *Pg* at an MOI of 50 for an additional 3 h. The cells were then harvested using Accutase (Corning Incorporated) and fixed with 4% PFA for 15 min. After washing with PBS, the cell pellets were incubated in PBS with BODIPY 493/503 for 30 min at RT, protected from light. The cells were washed and resuspended in PBS and analyzed on Symphony flow cytometry (BD Bioscience). MFI of BODIPY 493/503 staining was analyzed using FlowJo software.

**In vitro analysis of reactive oxygen species (ROS)** **production in BV2 cells**

To investigate the effect of *Pg* on ROS production in BV2 cells, as well as the interactions between LD accumulation and ROS generation, BV2 cells were plated on 8-well chamber slides (for confocal microscopy analysis) or in 24-well plates (for flow cytometry analysis) as described above. Cells were pretreated with vehicle, TrC, or 500 μM of N-Acetylcysteine amide (AD4; Cyman Chemical), a ROS inhibitor, for 3 h, followed by infection with *Pg* (MOI = 50) for an additional 3 h. The cells were then incubated with CellROX^TM^ Deep Red (Thermo Fisher Scientific; 1:500 in culture medium) or BODIPY 493/503 for 30 min at 37°C. After staining, cells were processed for confocal microscopy or flow cytometry analysis as described above.

**In vitro analysis of BV2 cells phagocytosis of Aβ peptide**

To investigate the effect of *Pg* on the phagocytosis ability of BV2 cells, as well as the role of LD accumulation in regulating phagocytosis, BV2 cells were plated on 8-well chamber slides (for confocal microscopy analysis) or in 24-well plates (for flow cytometry analysis) as described above. Cells were pretreated with vehicle, TrC, or 10 μM cytochalasin D (CytoD; Cyman Chemical), a phagocytosis inhibitor, for 3 h, followed by infection with *Pg* (MOI = 50) for an additional 3 h, and then incubated with 1.0 μM of Aβ (1-42)-HiLyte^TM^ Flour 488 (AnaSpec Inc, CA, USA) for another 2 h. After staining, cells were processed for confocal microscopy or flow cytometry analysis as described above.

**RT-qPCR**

To analyze the expression of genes related to oxidative stress (iNOS and NOX-2) and microglia activation (CD68, MHC-II, TMEM119), BV2 cells were homogenized, and total RNA was extracted using Trizol reagent. cDNA was synthesized using the PrimeScript RT Reagent kit with an equal amount of RNA. Real-time PCR was performed, and gene expressions were normalized to GAPDH expression, as described above. All primers used in this study are listed in **Appendix table 1**.

**In vivo infection model and tissue preparation**

For *in vivo* *Pg* infection, age- and sex-matched *App* KI mice (n=18; 21-25g) were randomly divided into three groups (n=6 per group): a control group (PBS only), a *Pg*-infected group, and a *Pg*+TrC group. Mice were given vehicle or TrC (10 mg/kg) via intraperitoneal (ip) injection once per day for two consecutive days, followed by *Pg* infection (1x10^7^ cfu/ml; 100 µl) via retro-orbital injection (Gaddis et al. 2013). Following 6 h of *Pg* infection, groups of mice were anesthetized with an ip injection of ketamine/xylene solution and transcardially perfused with PBS (Hao et al. 2022). Following perfusion, brains were collected and fixed in 4% PFA for 48 h. The brains were then cryoprotected in 30% sucrose in PBS overnight, sectioned into 45 μm coronal sections using a cryostat and stored in cryoprotected solution (glycerol: ethylene glycol: PBS, 1:1:2, pH 7.4) at -20°C for further experiments. The data analysis was performed in a blinded manner using coded samples to minimize bias.

**Immunofluorescence analysis of Ab plaques and LD accumulation in microglia from *App* KI mice**

Coronal brain sections were first stained for Ab plaques and subsequently co-stained for LD and microglia. For Aβ plaque detection, the sections were incubated with methoxy-X04 (Cayman Chemicals; 10 μM in PBS) for 30 min. The sections were washed three times with PBS, followed by blocking with blocking buffer (4.0% normal donkey serum and 0.2% Triton X-100 in PBS) for 2 h at RT. Following blocking, sections were washed and incubated with LipidSpot^TM^488 (Biotium; 1:1000 in PBS) for 3 h in the dark at RT. After three further washes, sections were incubated overnight at - 4°C with anti-rabbit IBA1 primary antibody (Abcam; 1:1000 in blocking buffer). The next day, sections were washed five times with PBS and incubated with donkey anti-rabbit IgG (H+L) Alexa fluor^TM^ 647 secondary antibodies (Thermo Fisher Scientific; 1:1000 in blocking buffer) for 2 h at RT. After three final washes, sections were mounted onto microscope slides using Fluromount-G^TM^ mounting medium without DAPI. Z-stack images were acquired using a confocal microscope with a 20x objective. To quantify LD^+^, IBA1^+,^ and LipisSpot^+^ cells, six visual fields were randomly selected. Cells with IBA1^+^, LipisSpot^+^, and methoxy-X04^+^ co-colonization were assessed to identify LD-accumulating microglia associated with Aβ peptides. All images were processed and analyzed using ImageJ software.

**Flow cytometry analysis of LD accumulation and ROS generation in microglia from *App* KI mice**

Brains were collected from mice in each group after perfusion and homogenized in Hank’s Balanced Salt Solution (HBSS) without calcium and magnesium (Thermo Fisher Scientific) using a 70 µm cell strainer (Thermo Fisher Scientific). Brain mononuclear cells were isolated following Percoll (Thermo Fisher Scientific) gradient centrifugation (30%, 37%, 70%) (Aggarwal et al. 2025; Lee and Tansey 2013). Cells at the 70% - 37% Percoll layer interfaces were collected and resuspended in FACS buffer. To analyze LD accumulation as well as the activation state of microglia, the cells were pre-incubated with anti-FC receptor antibody (BioLegend #101302; 1:500 in FACS buffer) on ice for 20 min. After washing with PBS, the cells were stained with anti-CD11b (BioLegend #101243; 1:100 in FACS buffer), anti-CD45 (BioLegend #103116; 1:100 in FACS buffer), anti-MCH-II (BioLegend #107612; 1:100 in FACS buffer), and anti-TMEM119 (eBioscience #2970936; 1:100 in FACS buffer) on ice for 30 min. After washing, the cells were incubated with LIVE/DEAD™ Fixable Aqua Stain (Thermo Fisher Scientific; 1:1000 in PBS) on ice for 20 min. Then the cells were fixed in 4% PFA for 10 min at RT. After washing, cells were incubated with BODIPY 493/503 for 30 min at RT, washed, and resuspended in PBS for flow cytometry analysis. To detect ROS generation in microglia, brain mononuclear cells were stained with CellROX Deep Red following CD11b and CD45 staining.

**Statistical analysis**

Statistical analyses were done using GraphPad Prism 10.0 software. All values were expressed as the means ± standard deviation, unless otherwise stated. Student *t*-test was used to determine the significance between the two groups. Comparisons between more than two groups were analyzed by the one-way analysis of variance (ANOVA). A *P* value < 0.05 was considered as a measure of statistical significance, with the different significance levels denoted as follows: * *P* < 0.05, ** *P* < 0.01, *** *P* < 0.00, and **** *P* < 0.0001.

**Appendix table 1:** Primer for RT-qPCR

| Primer | Forward | Reverse |
| --- | --- | --- |
| MCH-II | ACCCAGCCAAGATCAAAGTGC | TGCTCCACGTGACAGGTGTAGA |
| TMEM119 | CCTACTCTGTGTCACTCCCG | CACGTACTGCCGGAAATC |
| CD68 | TGTCTGATCTTGCTAGGACCG | GAGAGTAACCTTTTTGTGA |
| iNOS | GTTCTCAGCCCAACAATACAAGA | GTGGACGGGTCGATGTCAC |
| NOX-2 | TGTGGTTGGGGCTGAATGTC | CTGAGAAAGGAGAGCAGATTTCG |
| 16s rRNA | CTTGACTTCAGTGGCGGCA | AGGGAAGACGGTTTTCACCA |

**Appendix figures**

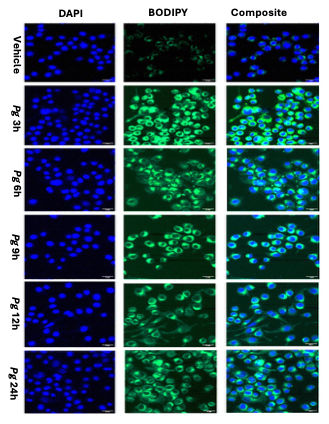

**Appendix figure 1: *Pg* infection accelerates the lipid droplet formation in BV2 microglia cells in a time-dependent manner**. The BV2 microglia cells were treated with vehicle or *Pg* (MOI=50) for 3h, 6h, 9h, 12h, and 24h. Representative confocal microscopy images of BV2 microglia cells primarily stained with BODIPY 493/503 (green) and counterstained with DAPI (blue); scale bar 20µm.

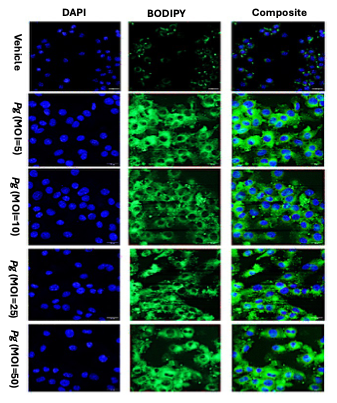

**Appendix figure 2: *Pg*-infection promotes LD accumulation in BV2 cells in a dose-dependent manner.** BV2 microglial cells were treated with *Pg* at MOI of 5, 10, 25, and 50 for 3h. The LD were stained with BODIPY 493/503 (green), and nuclei were counterstained with DAPI (blue); scale bar 20µm.

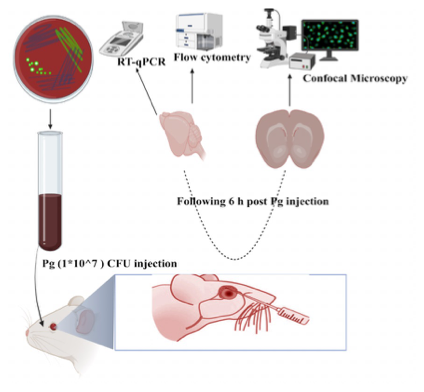

**Appendix figure 3: *Pg* retro-orbitally injection and downstream processing of samples.** Mice were retro-orbitally injected with *Pg (1 × 10^7 CFU)* and brains were collected 6h post-infection. Brain samples were processed for confocal microscopy (Zeiss LSM-800 Airy scan) to visualize microglial lipid droplet accumulation, flow cytometry (FACSymphony A5) to quantify microglial activation and functional status, and RT-qPCR analysis to evaluate expression of lipid metabolism, oxidative stress, and microglial activation status-related genes.

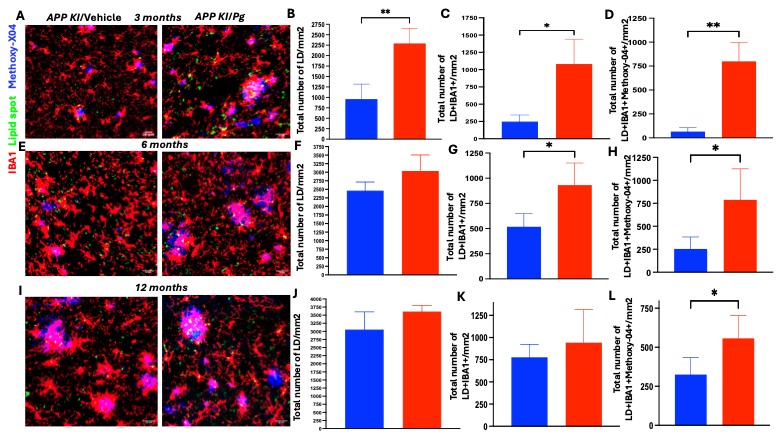

**Appendix figure 4:** **Pg infection exacerbates the microglia hypertrophy and aggregation in *App KI* mice**. **(A-D)** Representative confocal image (Scale bars, 20µm) and quantification of total LD, LD-loaded microglia, and LD-loaded microglia in Aβ positive area from the brain of 3-month-old App KI mice. **(E-H)** Representative confocal image (Scale bars, 20µm) and quantification of total LD, LD-loaded microglia, and LD-loaded microglia in Aβ positive area from the brain of 6-month-old App KI mice. **(I-L)** Representative confocal image (Scale bars, 20µm) and quantification of total LD, LD-loaded microglia, and LD-loaded microglia in Aβ positive area from the brain of 12-month-old App KI mice. The data are presented as the mean ± SEM. P<0.0001 (n =6). *, *P* < 0.05, ** *P* < 0.01, *** *P* < 0.001 by Welch’s t test.

**References**

Aggarwal A, Mendoza-Mari Y, Aggarwal A, Agrawal DK. 2025. Isolation of primary brain cells: Challenges and solutions. Arch Clin Biomed Res. 9(4):286-296.

Gaddis DE, Maynard CL, Weaver CT, Michalek SM, Katz J. 2013. Role of tlr2-dependent il-10 production in the inhibition of the initial ifn-γ t cell response to *Porphyromonas gingivalis*. J Leukoc Biol. 93(1):21-31.

Hao X, Li Z, Li W, Katz J, Michalek SM, Barnum SR, Pozzo-Miller L, Saito T, Saido TC, Wang Q et al. 2022. Periodontal infection aggravates c1q-mediated microglial activation and synapse pruning in alzheimer’s mice. Front Immunol. Volume 13 - 2022.

Lee JK, Tansey MG. 2013. Microglia isolation from adult mouse brain. Methods Mol Biol. 1041:17-23.

Saito T, Matsuba Y, Mihira N, Takano J, Nilsson P, Itohara S, Iwata N, Saido TC. 2014. Single app knock-in mouse models of alzheimer's disease. Nat Neurosci. 17(5):661-663.

Yazdanpanah Moghadam E, Sonenberg N, Packirisamy M. 2025. Alzheimer model chip with microglia bv2 cells. Microsyst Nanoeng. 11(1):135.

Zhao Y, Li Z, Su L, Ballesteros-Tato A, Katz J, Michalek SM, Feng X, Zhang P. 2020. Frontline science: Characterization and regulation of osteoclast precursors following chronic *Porphyromonas gingivalis* infection. J Leukoc Biol. 108(4):1037-1050.
